## Supplementary Materials for "No effects of offline high frequency transcranial magnetic stimulation to posterior parietal cortex on the choice of which hand to use to perform a reaching task"

### S1. Supplementary statistical analyses

#### Table S1. Hand choice

(S1.1) Full dataset (N = 26)

***One-way ANOVA:***

*PSE per stimulation condition: $F(2, 50) = 1.73$, $p = 0.19$*
*PSE per stimulation condition with No-cTBS: $F(3, 75) = 1.58$, $p = 0.20$*

*Proportion RHU per stimulation condition: $F(2, 50) = 1.05$, $p = 0.36$*
*Proportion RHU per stimulation condition with No-cTBS: $F(3, 75) = 0.98$, $p = 0.41$*

*Proportion RHU at Sham-PSE per stimulation condition: $F(2, 50) = 1.23$, $p = 0.30$*
*Proportion RHU at No-cTBS-PSE per stimulation condition with No-cTBS: $F(3, 75) = 3.35$, $p = 0.02$*

(S1.2) Left-handers removed (N = 23)

***One-way ANOVA:***

*PSE per stimulation condition: $F(2, 44) = 1.44$, $p = 0.25$*
*PSE per stimulation condition with No-cTBS: $F(3, 66) = 1.32$, $p = 0.28$*

*Proportion RHU per stimulation condition: $F(2, 44) = 0.97$, $p = 0.39$*
*Proportion RHU per stimulation condition with No-cTBS: $F(3, 66) = 0.91$, $p = 0.44$*

*Proportion RHU at Sham-PSE per stimulation condition: $F(2, 44) = 1.41$, $p = 0.26$*
*Proportion RHU at No-cTBS-PSE per stimulation condition with No-cTBS: $F(3, 66) = 3.38$, $p = 0.02$*

(S1.3) Right-handers with strategy removed (N = 24)

***One-way ANOVA:***

*PSE per stimulation condition: $F(2, 46) = 1.57$, $p = 0.22$*
*PSE per stimulation condition with No-cTBS: $F(3, 69) = 1.40$, $p = 0.25$*

*Proportion RHU per stimulation condition: $F(2, 46) = 0.99$, $p = 0.38$*
*Proportion RHU per stimulation condition with No-cTBS: $F(3, 69) = 0.94$, $p = 0.43$*

*Proportion RHU at Sham-PSE per stimulation condition: $F(2, 46) = 0.99$, $p = 0.38$*
*Proportion RHU at No-cTBS-PSE per stimulation condition with No-cTBS: $F(3, 69) = 3.07$, $p = 0.03$*

(S1.4) TMS-averse removed (N = 25)

***One-way ANOVA:***

*PSE per stimulation condition: $F(2, 48) = 1.63$, $p = 0.21$*
*PSE per stimulation condition with No-cTBS: $F(3, 72) = 1.50$, $p = 0.22$*

*Proportion RHU per stimulation condition: $F(2, 48) = 0.99$, $p = 0.38$*
*Proportion RHU per stimulation condition with No-cTBS: $F(3, 72) = 0.92$, $p = 0.43$*

*Proportion RHU at Sham-PSE per stimulation condition: $F(2, 48) = 1.18$, $p = 0.32$*
*Proportion RHU at No-cTBS-PSE per stimulation condition with No-cTBS: $F(3, 72) = 3.20$, $p = 0.03$*

(S1.5) Right-handers, no strategy (N = 20)

***One-way ANOVA:***

*PSE per stimulation condition: $F(2, 38) = 1.19$, $p = 0.31$*
*PSE per stimulation condition with No-cTBS: $F(3, 57) = 1.07$, $p = 0.37$*

*Proportion RHU per stimulation condition: $F(2, 38) = 0.84$, $p = 0.44$*
*Proportion RHU per stimulation condition with No-cTBS: $F(3, 57) = 0.80$, $p = 0.50$*

*Proportion RHU at Sham-PSE per stimulation condition: F (2, 38) = 1.09, p = 0.35*
*Proportion RHU at No-cTBS-PSE per stimulation condition with No-cTBS: F (3, 57) = 2.96, p = 0.04*

(S1.6) Right-handers, no strategy, outlier removed (N = 19)

***One-way ANOVA:***

*PSE per stimulation condition: F (2, 36) = 0.56, p = 0.58*
*PSE per stimulation condition with No-cTBS: F (3, 54) = 0.48, p = 0.70*

*Proportion RHU per stimulation condition: F (2, 36) = 0.71, p = 0.50*
*Proportion RHU per stimulation condition with No-cTBS: F (3, 54) = 0.60, p = 0.62*

*Proportion RHU at Sham-PSE per stimulation condition: F (2, 36) = 1.26, p = 0.30*
*Proportion RHU at No-cTBS-PSE per stimulation condition with No-cTBS: F (3, 54) = 2.51, p = 0.07*

### Table S2. Response times

(S2.1) Full dataset (N = 26)

***Two-way ANOVA:*** **Hand x Stimulation condition**

*Main effect of Hand: F (1, 25) = 1.83, p = 0.19*
*Main effect of Stimulation condition: F (2, 50) = 0.19, p = 0.83*
*Interaction term: F (2, 50) = 0.38, p = 0.69*

***Choice costs***

**Space x Stimulation condition: Sham-PSE**
*Main effect of Space: F (1, 25) = 18.33, p = 0.0002*
*Main effect of Stimulation condition: F (2, 50) = 0.11, p = 0.89*
*Interaction term: F (2, 50) = 1.40, p = 0.26*

**Space x Stimulation condition: No-cTBS-PSE**
*Main effect of Space: F (1, 25) = 16.51, p = 0.0004*
*Main effect of Stimulation condition: F (2, 50) = 0.18, p = 0.84*
*Interaction term: F (2, 50) = 1.42, p = 0.25*

(S2.2) Left-handers removed (N = 23)

***Two-way ANOVA:*** **Hand x Stimulation condition**

*Main effect of Hand: F (1, 22) = 0.82, p = 0.38*
*Main effect of Stimulation condition: F (2, 44) = 0.86, p = 0.43*
*Interaction term: F (2, 44) = 0.10, p = 0.90*

***Choice costs***

**Space x Stimulation condition: Sham-PSE**
*Main effect of Space: F (1, 22) = 15.10, p = 0.0008*
*Main effect of Stimulation condition: F (2, 44) = 0.66, p = 0.52*
*Interaction term: F (2, 44) = 1.82, p = 0.17*

**Space x Stimulation condition: No-cTBS-PSE**
*Main effect of Space: F (1, 22) = 14.95, p = 0.0008*
*Main effect of Stimulation condition: F (2, 44) = 0.33, p = 0.72*
*Interaction term: F (2, 44) = 0.69, p = 0.51*

(S2.3) Right-handers with strategy removed (N = 24)

***Two-way ANOVA:*** **Hand x Stimulation condition**

*Main effect of Hand: F (1, 23) = 2.50, p = 0.13*
*Main effect of Stimulation condition: F (2, 46) = 0.21, p = 0.81*
*Interaction term: F (2, 46) = 0.18, p = 0.84*

***Choice costs***

**Space x Stimulation condition: Sham-PSE**
*Main effect of Space: F (1, 23) = 15.68, p = 0.0006*
*Main effect of Stimulation condition: F (2, 46) = 0.15, p = 0.86*
*Interaction term: F (2, 46) = 0.87, p = 0.43*

**Space x Stimulation condition: No-cTBS-PSE**
*Main effect of Space: F (1, 23) = 13.96, p = 0.001*
*Main effect of Stimulation condition: F (2, 46) = 0.19, p = 0.83*
*Interaction term: F (2, 46) = 0.90, p = 0.41*

(S2.4) TMS-averse removed (N = 25)

***Two-way ANOVA:*** **Hand x Stimulation condition**

*Main effect of Hand: F (1, 24) = 1.48, p = 0.24*
*Main effect of Stimulation condition: F (2, 48) = 0.13, p = 0.88*
*Interaction term: F (2, 48) = 0.36, p = 0.70*

***Choice costs***

**Space x Stimulation condition: Sham-PSE**
*Main effect of Space: F (1, 24) = 16.71, p = 0.0004*
*Main effect of Stimulation condition: F (2, 48) = 0.06, p = 0.94*

*Interaction term: F (2, 48) = 1.51, p = 0.23*

**Space x Stimulation condition: No-cTBS-PSE**
*Main effect of Space: F (1, 24) = 14.99, p = 0.0007*
*Main effect of Stimulation condition: F (2, 48) = 0.11, p = 0.90*
*Interaction term: F (2, 48) = 1.62, p = 0.21*

(S2.5) Right-handers, no strategy (N = 20)

***Two-way ANOVA:*** **Hand x Stimulation condition**
*Main effect of Hand: F (1, 19) = 1.05, p = 0.32*
*Main effect of Stimulation condition: F (2, 38) = 0.85, p = 0.43*
*Interaction term: F (2, 38) = 0.07, p = 0.93*

***Choice costs***
**Space x Stimulation condition: Sham-PSE**
*Main effect of Space: F (1, 19) = 11.02, p = 0.004*
*Main effect of Stimulation condition: F (2, 38) = 0.64, p = 0.53*
*Interaction term: F (2, 50) = 1.24, p = 0.30*

**Space x Stimulation condition: No-cTBS-PSE**
*Main effect of Space: F (1, 19) = 12.90, p = 0.002*
*Main effect of Stimulation condition: F (2, 38) = 0.71, p = 0.50*
*Interaction term: F (2, 50) = 0.82, p = 0.45*

(S2.6) Right-handers, no strategy, outlier removed (N = 19)

***Two-way ANOVA:*** **Hand x Stimulation condition**
*Main effect of Hand: F (1, 18) = 0.73, p = 0.40*
*Main effect of Stimulation condition: F (2, 36) = 0.90, p = 0.42*
*Interaction term: F (2, 36) = 0.41, p = 0.66*

***Choice costs***
**Space x Stimulation condition: Sham-PSE**
*Main effect of Space: F (1, 18) = 9.05, p = 0.008*
*Main effect of Stimulation condition: F (2, 36) = 0.72, p = 0.49*
*Interaction term: F (2, 36) = 0.67, p = 0.52*

**Space x Stimulation condition: No-cTBS-PSE**
*Main effect of Space: F (1, 18) = 10.75, p = 0.004*
*Main effect of Stimulation condition: F (2, 36) = 0.79, p = 0.46*
*Interaction term: F (2, 36) = 0.26, p = 0.77*

### S3. Participant errors

*Table S3. Participant errors.*

| Participant | Total | Percent of Participant's trials | Negative response time | Double button response |
|---|---|---|---|---|
| **1** | 2 | 0.10 | 0 | 2 |
| **2** | 15 | 0.75 | 3 | 13 |
| **3** | 10 | 0.50 | 9 | 1 |
| **4** | 6 | 0.30 | 2 | 4 |
| **5** | 12 | 0.59 | 3 | 9 |
| **6** | 7 | 0.35 | 4 | 3 |
| **7** | 14 | 0.69 | 4 | 10 |
| **8** | 17 | 0.84 | 13 | 4 |
| **9** | 13 | 0.64 | 6 | 7 |
| **10** | 15 | 0.74 | 4 | 11 |
| **11** | 6 | 0.30 | 0 | 6 |
| **12** | 12 | 0.60 | 7 | 5 |
| **13** | 13 | 0.64 | 7 | 6 |
| **14** | 1 | 0.05 | 1 | 0 |
| **15** | 22 | 1.09 | 12 | 10 |
| **16** | 7 | 0.35 | 2 | 5 |
| **17** | 6 | 0.30 | 3 | 3 |
| **18** | 7 | 0.35 | 4 | 3 |
| **19** | 9 | 0.45 | 7 | 2 |
| **20** | 11 | 0.55 | 7 | 4 |

**S4. Additional analyses: No-cTBS baseline**

We decided to perform a separate set of exploratory analyses using No-cTBS as the baseline measure of hand choice, rather than Sham-cTBS. Our motivations for these additional analyses were twofold. First, we wanted to address the possibility that Sham-cTBS may have influenced hand choice, perhaps by changing participant expectations—i.e., placebo effects. Some participants reported experiencing Sham-cTBS as real stimulation, and reported post-stimulation-related sensations following Sham-cTBS (see Supplementary Materials S7). If Sham-cTBS inadvertently influenced subsequent hand choice behaviour this may have obscured our ability to detect differences between conditions; at least in principle (for e.g., if all conditions were to shift hand choice in a similar direction). Second, the primary TMS results reported by Oliveira et al. (1) involved comparison with a No-TMS baseline. This motivates us to also consider comparison of real cTBS to a No-cTBS baseline.

The same three sets of tests used in our main preregistered analyses were performed, yet with four conditions included: (1) No-cTBS; (2) Sham-cTBS; (3) L-pIP-SPC; (4) R-pIP-SPC. No-cTBS was defined by the combined pre-stimulation data from Sessions 2 and 3. Pre-stimulation data from Session 1 were considered practice. These analyses were not preregistered.

First, we examined PSE measures. The results reveal no significant differences is PSEs between conditions ($F(3, 54) = 0.48$, $p = 0.70$, $\eta^2_p = 0.03$) (Figure S4.1A). The group mean PSEs for No-cTBS and Sham-cTBS are similar (-6.7˚ and -7˚, respectively). These results suggest that the use of Sham-cTBS as a baseline measure of hand choice is uncomplicated by placebo effects, at least at the group-level.

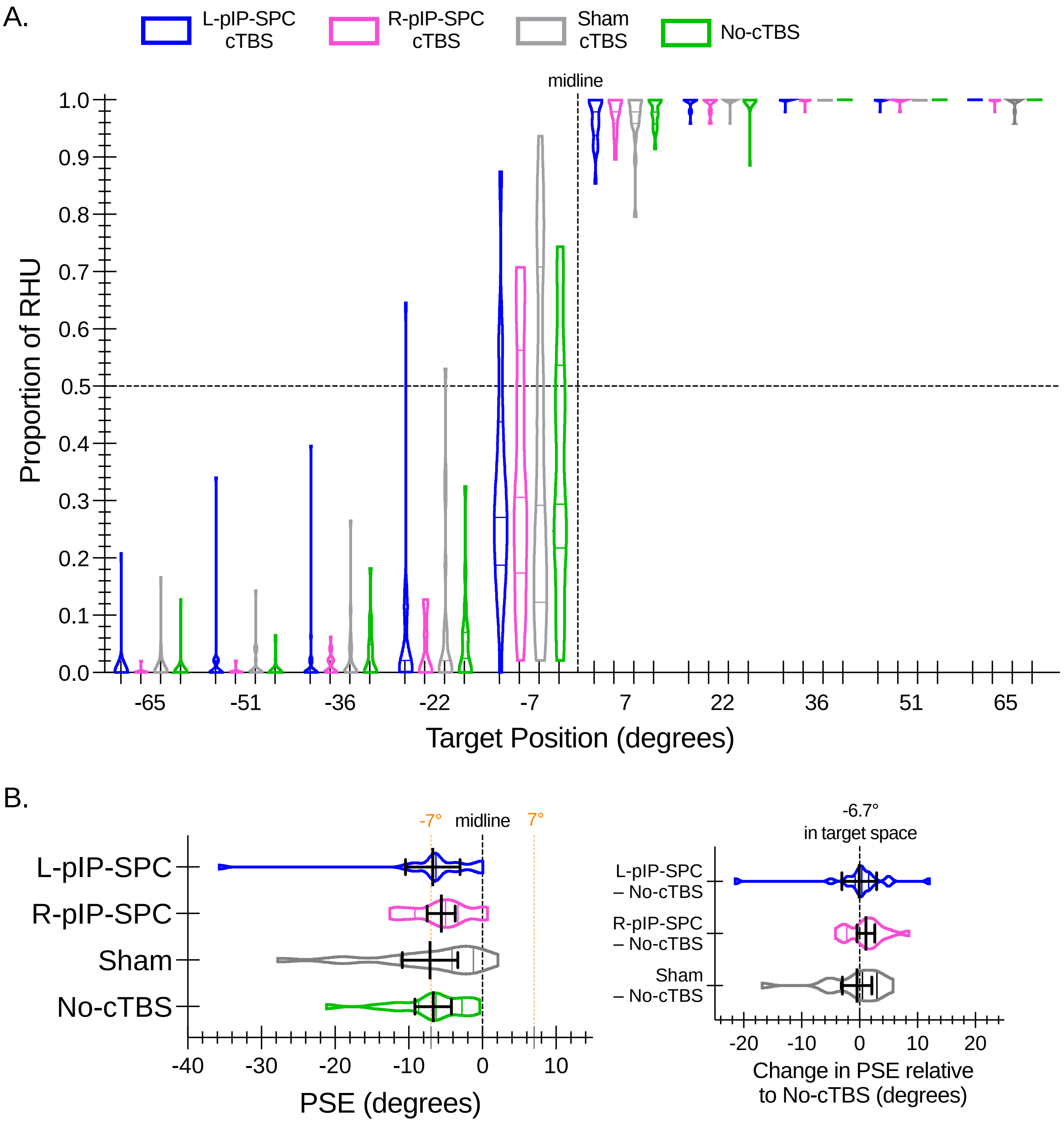


**Figure S4.1. Hand choice: No-cTBS. (A)** Violin plots show the interparticipant distribution of hand choice data across target locations expressed as the proportions of right-hand use (RHU) for L-pIP-SPC (blue), R-pIP-SPC (pink), Sham (grey), and No-cTBS (green) stimulation conditions. Within each violin plot the median and upper and lower quartile values are indicated. A vertical dashed line depicts the midline of the display (0°). A horizontal dashed line shows the point of equal proportion of left- and right-hand use. **(B)** Violin plots show the interparticipant distribution of mean PSEs per condition. Solid black lines indicate group means with 95% confidence intervals. The locations of the midline and targets -7° and 7° are shown for reference. *Inset*. Difference scores of PSEs for each condition relative to No-cTBS are shown as violin plots. Group means and 95% confidence intervals are overlaid.

Second, we examined the arcsine transformed proportions of RHU across all target locations. These results also reveal no significant differences between conditions $F(3, 54) = 0.60$, $p = 0.62$, $\eta^2_p = 0.03$) (Figure S4.2A).

Lastly, focusing on those targets that bound the PSE, the results reveal significant effects of cTBS ($F(3, 57) = 2.96$, $p < 0.05$, $\eta^2_p = 0.14$) (Figure S4.2B). Follow-up tests show that these results reflect significant differences between R-pIP-SPC and Sham-cTBS conditions. The likelihood of right-hand choice is decreased after cTBS to the R-pIP-SPC relative to Sham-cTBS. This pattern is evident when compared with No-cTBS, yet these differences do not survive correction for multiple comparisons.

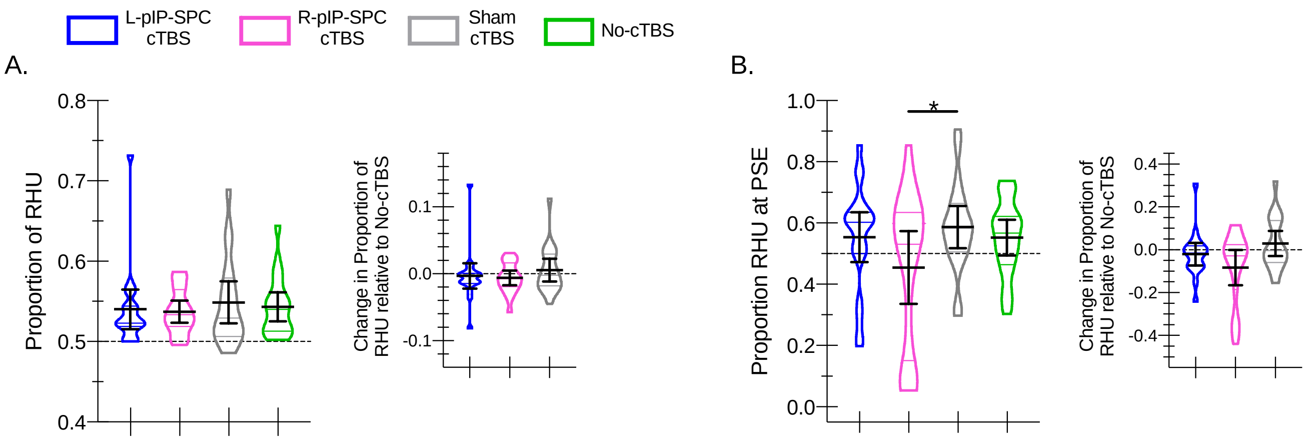


**Figure S4.2. Hand choice: Proportion of right hand use: No-cTBS. (A)** Violin plots depict the distribution of hand choice data collapsed across targets expressed as the proportions of right-hand use (RHU) for L-pIP-SPC (blue), R-pIP-SPC (pink), Sham (grey), and No-cTBS (green) stimulation conditions. Within each violin plot the median and upper and lower quartile values are indicated. Solid black lines indicate group means with 95% confidence intervals. *Inset.* Difference scores show the proportion of RHU per condition relative to No-cTBS. Data are shown as violin plots with group means and 95% confidence intervals overlaid. **(B)** Same as in **(A)** yet restricted to those targets that bound the PSE.

Inspection of individual-level data shows that the differences in outcomes between these exploratory and our preregistered analyses reflect a change in PSE estimates, and thus corresponding PSE-bounding targets for six (of 20) participants. In all cases, the PSE-bounding targets are positively shifted (i.e., rightwardly) in target space when defined by No-cTBS compared to Sham-cTBS. In five participants the PSE shifts from between targets -22˚ and -7˚ as defined by Sham-cTBS to targets -7˚ and

+7˚ as defined by No-cTBS. The sixth participant shows a shift from targets -36˚ and -22˚, Sham-cTBS, to targets -22˚ and -7˚, No-cTBS. The pattern is consistent with a tendency for Sham-cTBS to cause a relative increase in right-hand choice, if we assume that No-cTBS reflects a ‘true' baseline. Altogether these findings are difficult to interpret. At the group-level, we observe highly variable hand choice behaviour following Sham-cTBS.

To summarise, our exploratory results reveal subtle changes in hand choice following cTBS; in particular, the probability of right-hand choice is reduced following cTBS to the right hemisphere pIP-SPC. The effects are restricted to the area of target space where participants are most likely to use either hand, and show considerable interparticipant variability. Other analyses fail to yield statistically significant outcomes, although the same pattern of comparatively reduced right-hand choice following cTBS to R-pIP-SPC is observed. We offer these data as preliminary evidence that may serve of value for future studies, and that, until tested further, should be interpreted cautiously.

We also repeated our RT analyses with No-cTBS as the baseline. The statistical outcomes are the same as those reported in the main manuscript (Section 3.2), using Sham cTBS as baseline.

### S5. Comparison of reaching studies

A synthesis of reported prior neuroimaging and TMS results involving reaching and pointing is presented in Figure 5 of the paper. Specifically, we used the following method to provide an objective estimate of the overlap in reported effects across studies. We take the reported coordinates of peak activation (fMRI/PET) or target area (TMS coil localisation) and create a 15mm-diameter spherical foci of ‘activity’ around these coordinates, as the centroid. We then use these projected ‘activations’ to compute a heat map representing the percent overlap between studies, projected on the three-dimensional cortical surface of a single individual in standardised (Talairach) space. Foci with a minimum of 20% overlap with other coordinates are illustrated. Our choice of 15mm-diameter foci with ≥20% overlap to represent the variability, or noise, in the location of these reported coordinates was otherwise arbitrary. A total of 14 fMRI, three PET, and 7 TMS studies were included. Where necessary, reported MNI coordinates were transformed to Talairach coordinates (5), using the approach provided by the Cambridge Brain Sciences Unit (6). This provides a simple way to visualise the variation/consistency in reported functional-localisation data from prior studies involving reaching and arm movements alongside the coordinates we used as a guide for TMS coil localisation in the current study, based on our own recent fMRI results (7).

### S6. Post-stimulation questionnaire data

*Table S6.1.* Post-stimulation questionnaire data. Full dataset (N = 26)

**A.**

| Type of Session | Reported as Real | Reported as Sham | Reported "I don't know" | Percent correct identification |
|---|---|---|---|---|
| | | | | *Total sample* \| *Respondents* |
| **Real cTBS** | 28 | 5 | 19 | 53.85% \| 84.85% |
| **Sham cTBS** | 13 | 2 | 11 | 7.69% \| 13.33% |

**B.**

| Type of Sensation | Reported after Real | Reported after Sham |
|---|---|---|
| **Itching** | 6 | 1 |
| **Pain** | 6 | 1 |
| **Burning** | - | 1 |
| **Warmth/heat** | 7 | 4 |
| **Pinching** | 6 | 1 |
| **Metallic/iron taste** | 2 | - |
| **Fatigue** | 8 | 4 |
| **Dazed** | 1 | - |
| **Tapping** | 4 | 1 |
| **Twitching** | 1 | - |

*Table S6.2.* Post-stimulation questionnaire data. TMS-averse removed (N = 25)

**A.**

| Type of Session | Reported as Real | Reported as Sham | Reported “I don’t know” | Percent correct identification |
|---|---|---|---|---|
| | | | | *Total sample* \| *Respondents* |
| **Real cTBS** | 26 | 5 | 19 | 52.00% \| 81.25% |
| **Sham cTBS** | 12 | 2 | 11 | 8.00% \| 14.29% |

**B.**

| Type of Sensation | Reported after Real | Reported after Sham |
|---|---|---|
| **Itching** | 5 | 1 |
| **Pain** | 4 | - |
| **Burning** | - | - |
| **Warmth/heat** | 5 | 3 |
| **Pinching** | 5 | - |
| **Metallic/iron taste** | - | - |
| **Fatigue** | 6 | 3 |
| **Dazed** | 1 | - |
| **Tapping** | 4 | 1 |
| **Twitching** | 1 | - |

*Table S6.3.* Post-stimulation questionnaire data. Right-handers, no strategy (N = 20)

**A.**

| **Type of Session** | **Reported as Real** | **Reported as Sham** | **Reported "I don't know"** | **Percent correct identification** |
|---|---|---|---|---|
| | | | | *Total sample* \| *Respondents* |
| **Real cTBS** | 19 | 4 | 17 | 47.50% \| 82.61% |
| **Sham cTBS** | 10 | 0 | 10 | 0% |

**B.**

| **Type of Sensation** | **Reported after Real** | **Reported after Sham** |
|---|---|---|
| **Itching** | 5 | 1 |
| **Pain** | 1 | - |
| **Burning** | - | - |
| **Warmth/heat** | 3 | 2 |
| **Pinching** | 4 | - |
| **Metallic/iron taste** | - | - |
| **Fatigue** | 4 | 1 |
| **Dazed** | 1 | - |
| **Tapping** | 4 | 1 |
| **Twitching** | 5 | 1 |
